# Natural killer cell function is regulated by TGF-β signaling in pregnancy and tumor progression

**DOI:** 10.1101/2024.09.18.613652

**Authors:** Adam Yalin, Shuang-Yin Wang, Tomer Landsberger, Moriya Gamliel, Naama Elefant, Rebecca Kotzur, Debra Goldman-Wohl, Eyal David, Martina Molgora, Bishan Bhattarai, Marina Cella, Tomer Meir-Salame, Ronit Gilad, Rana Abdelkader, David Shveiky, Simcha Yagel, Or Zuk, Marco Colonna, Ofer Mandelboim, Ido Amit

## Abstract

The immune compartment of the maternal-fetal interface must balance between supporting maternal-fetal interactions and maintaining maternal tolerance. Despite recent advances, the cellular and molecular regulators that drive maternal immune cell remodeling remain largely unknown. Using index and transcriptional single-cell sorting, we comprehensively characterized the immune compartment dynamics in the maternal-fetal interface of both human and mouse and charted the markers and functional pathways associated with these cells. Comparing immune signatures in decidua and tumors of human and mice, we identify conserved gene modules that are activated in natural killer (NK) cells of both compartments, including TFG-β signaling. Genetic ablation and antibody blockade of the TGF-β pathway in NK cells resulted in enhanced anti-tumor immunity.

## Introduction

The maternal-fetal interface (MFI) integrates two semiallogeneic subjects. Major components of this interface are fetal trophoblast cells and the maternal decidua, which are complex heterogeneous structures that comprise the mature placenta. The decidua is comprised of stromal cells, blood vessels and immune cells that foster the safe development of the semiallogeneic fetus^1,2^. Immune cells populating this niche, such as natural killer (NK) cells and macrophages, play key roles in regulating placental growth and spiral artery remodeling^3,4,5,6^ and their absence was linked to pregnancy pathologies such as pre-eclampsia^7,8^. NK cells were also shown to have important role in fetal protection from infections and pathogens^9^. Other immune populations such as decidual T cells and dendritic cells (DCs) are present at the MFI^10,11^. Decidual DCs were shown to possess reduced antigen presentation capabilities along with dampened migratory capacity^12^. Furthermore, decidual CD8 T cells were shown to express inhibitory receptors such as programed cell death 1 (PD-1) and LAG3^13^. These T cells are also unresponsive to the non-classical HLA-G-expressing trophoblast cells^14^. These poor adaptive immunity features at the MFI have an important role in facilitating maternal tolerance. T-regulatory cells (Tregs) have been shown to ensure immune tolerance towards the semiallogeneic fetus by inhibiting alloreactive T cells thorough the secretion of immune modulatory cytokines such as IL-10 and transforming growth factor-β (TGF-β)^15^. Moreover, Treg deficiency in mice and human is manifested by systemic immunopathology and may result in reproduction malfunctions^16,17^. Tregs were also shown to play a crucial role in tumor progression, and their presence at the tumor microenvironment (TME) is correlated with dampened immunotherapy response and poor prognosis^18^.

Indeed, some immune modulatory mechanisms that support pregnancy are also exploited by malignancies^19^. The MFI shares many similarities in cell types, signaling and microenvironment with cancerous processes, including immune modulation, angiogenesis, hypoxia and neo-antigen presentation. These signals might in turn drive the immune compartment towards inhibitory state and contribute to the tumor immune evasion.

NK cells, similar to effector CD8 T cells, play an important role in anti-tumor immunity by recognizing and killing transformed or damaged cells^20,21,22^. NK cells are known to infiltrate a variety of human tumors and, in recent years, emerged as potential targets for immunotherapy^23,24,25^. However, high heterogeneity within the NK compartment is currently limiting our ability to isolate signals that might be targeted in order to use these effector cells clinically.

Previous studies have suggested that multiple signals and factors within the TME may drive the unique phenotype acquired by tumor-infiltrating immune cells, driving suppressive phenotypes that further support tumor progression. In the context of NK cells, the suppressive cytokine transforming growth factor-betta (TGF-β) was suggested to play a major role in controlling NK cells functions in a variety of tissues and pathologies, including cancer. Moreover, it is well established that high levels of TGF-β can be found in both TME and MFI (ref), suggesting a shared mechanism.

Recent advancements in high-throughput genomic technologies have provided a more complete view of the MFI^26,27,28^ and the TME^29,30,31^. Single-cell RNA sequencing (scRNA-seq) allows an unbiased characterization of heterogeneous cell types and contributed enormously to our understanding of cellular networks and complexed tissues such as the lung and the adipose tissue^32,33^. In a past study, we used scRNA-seq to generate the first multi-tissue, homeostasis mouse NK cells atlas^34^, revealing tissue-specific imprinting on NK cells. Mapping intra-tumor NK compartment in this resolution may reveal immunotherapeutic targets that will allow us to unleash the anti-tumor potential of NK cells.

The trade-off between fetal tolerance through immune inhibition and maternal as well as fetal protection has been a focus of research for decades; yet, the cell types and the microenvironment supporting this trade-off remain only partially understood. Uncovering the underlying mechanisms, e.g., specialized cell types and signaling pathways that are activated in suppressive immune microenvironments, may contribute to better understanding of parallel mechanisms that are exploited by malignancies.

Here, we report an extensive characterization of the immune compartment within the maternal-fetal interface of early and mid-pregnancy in both human and mouse. Using massively parallel scRNA-seq and functional assays, we characterize the different immune cell types and the cellular kinetics throughout the first trimester of pregnancy. Within these populations, we demonstrate plasticity of decidual natural killer (dNK) cells and identify three distinct transcriptional modules directing different dNK functions in both human and mouse. We find parallel NK subsets in multiple human tumors and murine syngeneic tumor models, alongside tumor-specific imprinting. We identify TGF-β signaling as a major driver of NK cell programs in the MFI and TME. Blocking this pathway resulted in higher NK infiltration and superior anti-tumor activity.

## Results

### 1. Immune cell landscape of mouse maternal-fetal interface

Both human and murine MFI are composed of two major components: the maternal decidua and the fetal trophoblast-derived placenta. Although there are differences in the interface of the two species, they share central functional features of immune regulation. In order to cellularly and molecularly characterize these immune regulatory pathways, we sampled cells from both placenta and decidua at three time points during murine pregnancy: early first trimester, before placenta is well formed (E8.5); mid-point when the placenta is fully functional (E12.5); and a time point that parallels the shift between the human first and second trimesters (E14.5). To avoid contamination from fetal immune cells, which are also present in the placenta, we used a transgenic mouse reporter system where fetal cells are marked with a fluorescent marker (**Extended Data Fig. 1a-d, See methods**). For unbiased characterization of the immune compartment, we utilized a broad gating strategy including all immune cells except for neutrophils (CD45^+^, Ly6g^-^) **(Fig. 1a and Supplementary Table 1)**. Using MARS-seq^35^, we profiled a total of 14,145 quality controlled (QC)-passed maternal immune cells (**Extended Data Fig. 1e-f**). We applied the MetaCell algorithm^36^ to identify homogeneous and robust groups of cells (referred to as “metacells”; see Methods), resulting in a detailed map of 88 metacells organized into 8 broad immune cell types **(Fig. 1b)**. We classified the different groups based on expression levels of the most variable genes and used the differentially expressed genes to annotate broad myeloid and lymphoid cell types. The main groups of MFI immune cells included T cells (characterized by expression of *Cd3g*), B cells (*Cd79a*), NK cells (*Ncr1*), dendritic cells (DCs; *Cd209a*), plasmacytoid DCs (pDCs; *Siglech*), monocytes (*Lyz2*), macrophages (*Apoe*), and basophils (*Mcpt8*) (**Fig. 1c-d**). Finally, we detected a massive reorganization of the immune cell population in the placenta during pregnancy progression, with the most prominent changes observed in monocytes-macrophages replaced by DCs and by decidual NK cells replaced by T cells (**Fig. 1e-f**). Our analysis identified two major groups of dNKs, one of which expressed multiple chemokines including the DC-attracting chemokine *Xcl1*, while the other expressed high levels of granzymes (**Fig. 1c-d and Extended Data Fig. 2a**). the latter group is comprised from two subpopulations, one of which populating the early MFI (at E8.5) and replaced by the second group (**Fig. 1e-f)** expressing higher levels of granzymes, metalloproteinases (Mt1/2), Spp1 and the transcription factor Epas1 (Hif2a) which is the key regulator of hypoxia, indicating this subgroup is imprinted by the dynamic signals within the MFI (**Extended Data Fig. 2a**).

**Fig. 1.**
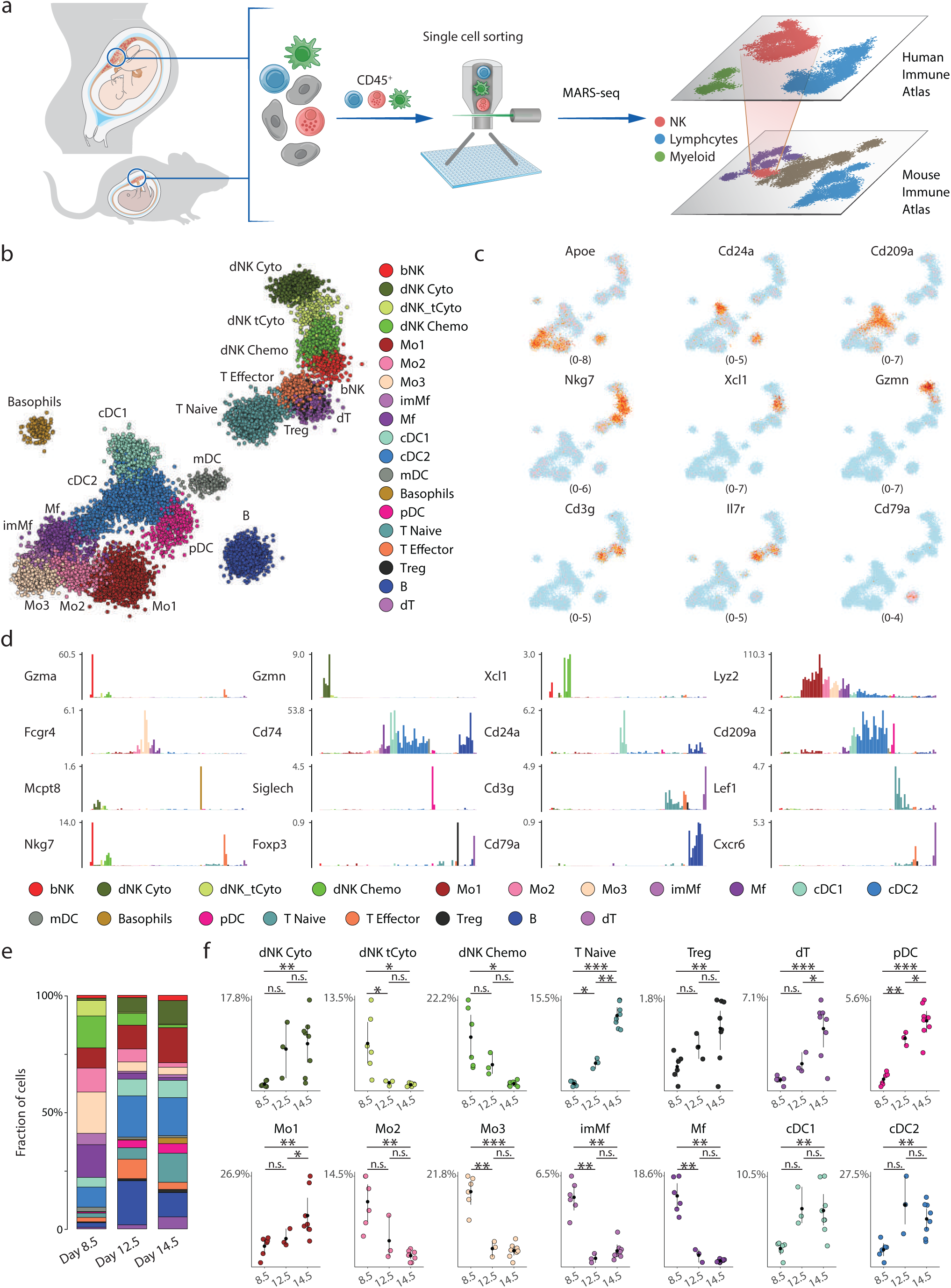
The immune cell landscape of mouse maternal-fetal interface across early to mid-pregnancy. **a,** Graphical overview of the experimental design. Using single cell sorting and our single-cell RNA-seq pipeline to study immune cells in mouse and human maternal-fetal interface tissues during pregnancy. **b,** Two-dimensional (2D) projection of transcriptome profiles of 14,145 QC-passed immune cells partitioned into 88 metacells. Single cells are taken from placentas of 9 mice from E8.5, E12.5, and E14.5 of pregnancy, and shown in dots. Different immune cell types are annotated and marked by color code. **c,** Heatmap showing unique molecular identifier (UMI) counts (log2 scale) of selected genes in individual cells on the 2D map in panel **b**. **d,** Normalized expression levels of select genes across the entire metacell model, color coded by cell type. **e**, Cell type compositions of the immune compartment in mouse MFI of E8.5, E12.5, and E14.5 of pregnancy. **f,** Frequency of different immune cell types within the three pregnancy time points (E8.5, E12.5, and E14.5) with each circle representing a placenta and error bars the time-point mean and standard deviation. Stars marking p value of a student t-test between time-point pairs (*p < 0.05; **p < 0.01; ***p < 0.001).

Similar to dNKs, decidual and uterine macrophages (dMafs) and monocytes displayed transcriptional dynamics along pregnancy progression (**Fig. 1e-f**). These dMafs were associated with anti-inflammation and tissue repair functions, as evident by the expression of genes such as *Ear2*, *Trem3*, and *Gngt2* (**Extended Data Fig. 2b**). We also detected two major dendritic cell subtypes; cDC1 (expressing *Naaa*, *Irf8*, and *Xcr1*) and cDC2 (expressing *Cd209a*) (**Fig. 1c-d**). Small decidual populations of pDCs and migratory DCs (expressing *Ccr7*, *Ccl17*, and *Ccl22*) were also observed. Lymphocytes were less abundant in the tissue. Whereas B cells and naïve T cells displayed little difference from circulating cells (**Extended Data Fig. 2c**), we found tissue resident Tregs (expressing *Foxp3*), decidual T cells (dT), both expressing the chemokine receptor *Cxcr6* and a small T cell population resembling effector CD8 T cells (expressing *Nkg7*, *Ccl5*, and *Itgb1*) **(Fig. 1c-d)**. In summary, our analysis revealed extensive changes in immune cell populations during pregnancy (**Extended data Fig. 2d**), including many tissue-resident populations that are unique to the MFI microenvironment.

**Fig. 2.**
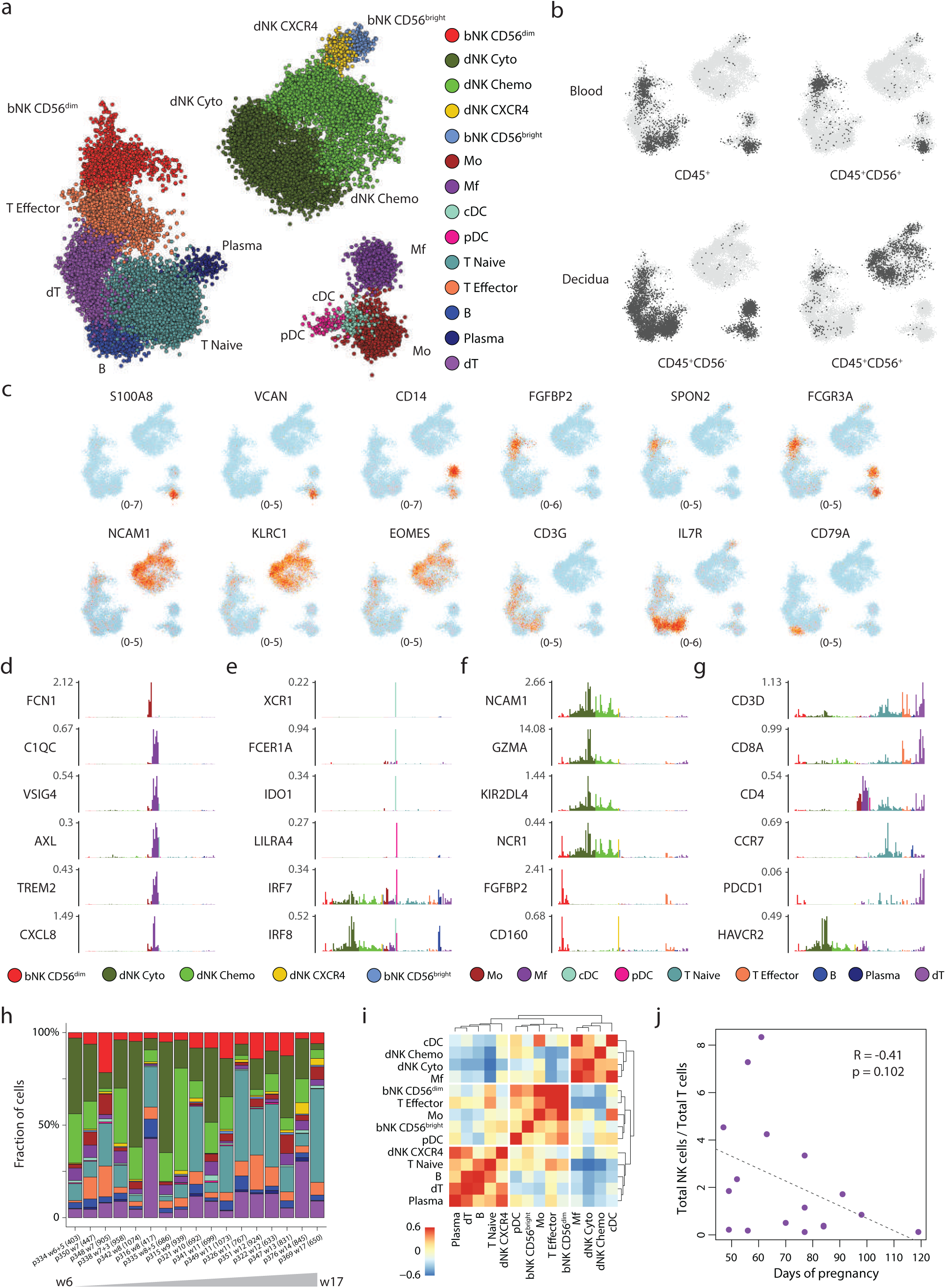
The immune cell landscape of human maternal-fetal interface across the first trimester. **a**, 2D projection of transcriptome profiles of 27,764 QC-passed immune cells partitioned into 125 metacells. Single cells are taken from 5 peripheral blood samples and 23 decidua tissue samples of elective first trimester termination of normal pregnancies. Individual cells and shown in dots. Different immune cell types are annotated and marked by color code. **b**, 2D projection of single cells by source (decidua or peripheral blood). Single cells from the indicated sample and gating are colored in dark grey on the 2D map in panel **a**. **c**, Heatmap showing unique molecular identifier (UMI) counts (log2 scale) of selected genes in individual cells on the 2D map in panel **a**. **d-g**, Normalized expression levels of select genes across the entire metacell model, color coded by cell type. **h**, Cell type compositions of the immune compartment in individual human decidual sample, ordered by the time point of pregnancy (by week) when the samples were taken. **i**, Heatmap showing the correlation of the frequency of different cell types within individual patients. **j**, The ratio of total NK cell to total T cells on the x axis, plotted against days of pregnancy on the y axis. Points designate individual patient samples. Dashed line is a linear regression line. R is the Pearson correlation coefficient and p is the p value for goodness of fit.

### 2. Immune cell landscape of human decidua

In order to compare between mouse and human immune landscapes, we collected 23 first trimester (6-14 weeks) decidua samples and 5 peripheral blood samples (**Extended Data Fig. 3a-b**). Using MARS-seq, we profiled a total of 27,764 QC-passed (**Extended data Fig. 3c-d**) maternal immune cells and applied the same analytical pipeline (**Fig. 2a and Supplementary Table 2**). In order to separate decidual resident from circulating immune cells, we compared decidual cells to immune cells isolated from peripheral blood (**Fig. 2b and Extended Data Fig. 3a-b**). Lymphoid cells constituted the majority of the human decidual immune cells. The most abundant myeloid cells were dMafs^5^ expressing *APOE*, complement genes (*C1QA*, *C1QB*, *C3*) and *TREM2*, which was shown to be expressed on several myeloid populations in the contexts of obesity and neurodegenerative diseases^37,32^ (**Fig. 2c-d**). Like their murine counterparts, dMafs expressed a large module of growth factors, cytokines and tissue residency genes, namely *CXCL8* (IL8), *CXCL2*/3, *IGF1*, *MRC1*, *CD163*, and *CD209* (*DC-SIGN*) (**Fig. 2d and Extended Data Fig. 4a**)^38,39,40^. They also expressed immunomodulatory genes such as *VSIG4* (**Fig. 2d**), a negative regulator of T-cell response. DCs (and pDCs) were relatively rare, in line with previous studies^10^ and expressed known DC markers *CD1C* and *XCR1* along with *IDO1*, a known modulator of T cell activity^41,42,43^ (**Fig. 2e and Extended Data Fig. 4a**).

**Fig. 3.**
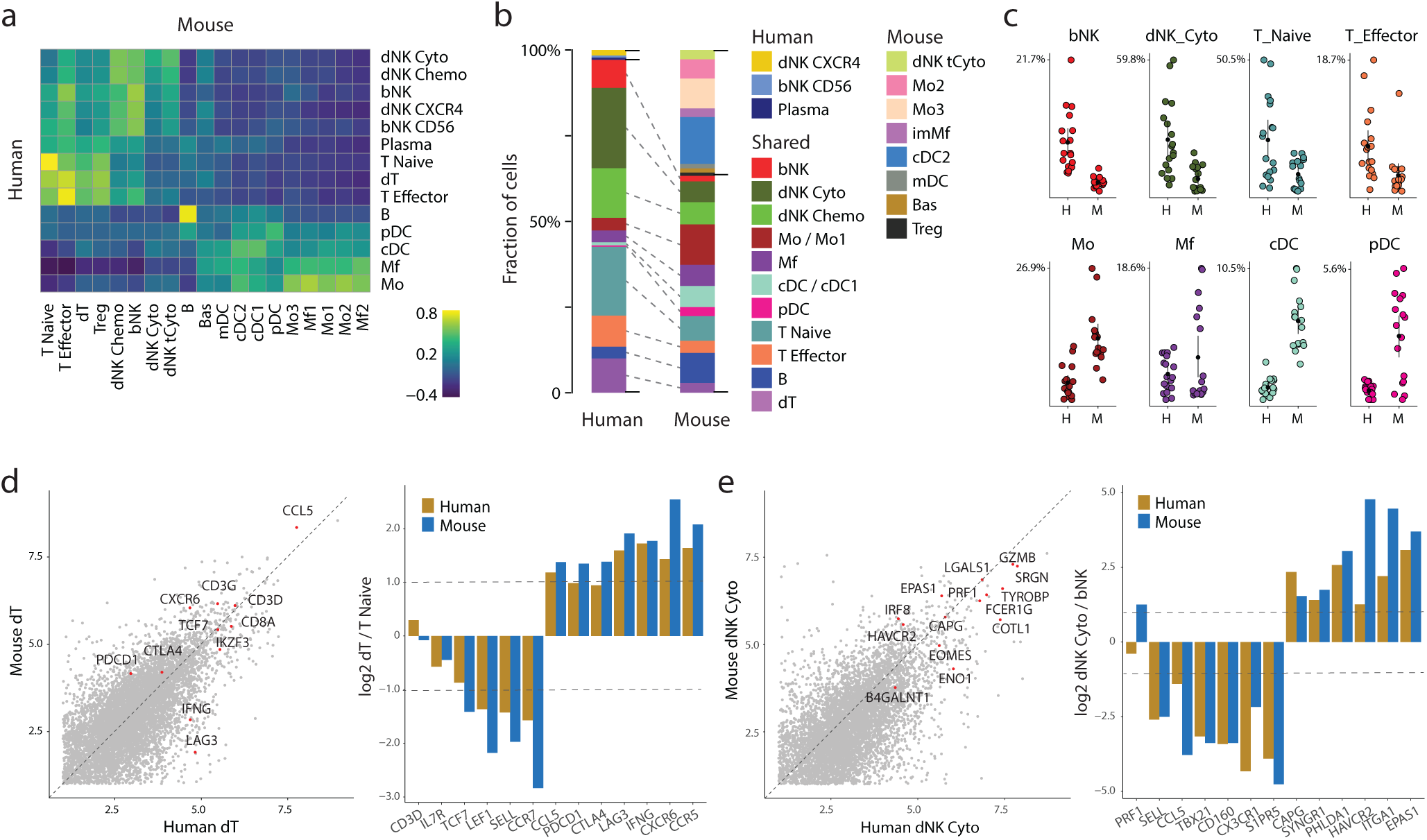
Comparison between the human and mouse maternal immune landscape across the first trimester. **a,** Heatmap showing similarities between individual human and mouse maternal immune cell types. **b**, Stacked barplot showing maternal immune cell type composition in human and mouse pregnancy. The same color codes are used for the immune cell types shared between human and mouse, while different color codes are used for the ones unique in human or mouse. **c**, Frequency of selected immune cell types within human and mouse pregnancy with each circle representing one sample. Lymphocyte and myeloid populations are shown in the top row and bottom row, respectively. **d**, Scatterplot on the left of the panel showing the average UMI counts (log2 scale) of human decidual T cells (x axis) compared with mouse decidual T cells (y axis), with the highlights of selected important genes. Barplot on the right of the panel showing expression fold change (log2 scale) of selected genes in human and mouse decidual T cells when compared with naïve T cells. **e**, same as panel **d**, but for cytotoxic decidua NK cells, and compared with circulating blood NK cells.

The largest lymphoid group contained decidual and blood NK cells. In line with the murine data, NK cells in the MFI were highly heterogeneous and included several molecular subtypes (**Fig. 2f**). A distinct decidual CD8^+^ T cell (dT) was also observed, which expressed high levels of T cell activation markers such as *CXCR6*, *CCR5*, *CCL5*, and *IFNG* (**Extended Data Fig. 4a**). dT also expressed many of the co-inhibitory checkpoint receptors such as *LAG3*, *TIGIT*, and *PDCD1* (**Fig. 2g and extended Data Fig. 4a**), a gene module that is mostly observed in cancer processes and chronic inflammation^44,45^, and potentially contributes to the tolerogenic properties of the immune compartment during pregnancy^13^. A smaller population (less than 1% of CD45+ cells) of decidual Tregs (*FOXP3*^+^, *CD25*^+^, *CTLA4*^+^) was identified. In recent years, Tregs have been studied extensively in the context of reproductive immunity^46,47^. These studies demonstrated their contribution in establishing and maintaining immune tolerance throughout pregnancy. According to these studies, Tregs are expanding from the point of fertilization, but their low abundance in our data suggests that they quickly diminish. Lastly, we found B cells and plasma cells in the decidua to be indistinguishable from their circulating counterparts (**Fig. 2b**), and likely originate from the massive network of blood vessels within the decidua.

Comparing the cellular dynamics during first trimester, we observed a greater fraction of dNK cells in the early stages of pregnancy (week 6-10), comprising up to 80% of the decidual immune cells, as compared to 5-30% after week 10 (**Fig. 2h and Extended Data Fig. 4b-c**), these results are in line with previous studies^48,49^. The reduction in dNK cells was compensated by an increase in T cells and peripheral blood CD56^dim^ CD16^+^ NK cells (**Fig. 2h-j**), possibly owing to changes in blood flow to the tissue. Yet, in later weeks some populations were highly decidua-specific, such as the dT cells, which showed major differences as compared to circulating T cells (**Fig. 2b**). We also observed correlated patterns in immune cellular networks and cell signaling. For example, in early pregnancy the fractions of dNKs and dMafs were correlated (R = 0.53, *p* = 0.024, **Fig. 2i**.). dNKs and dMafs may form cell-cell interactions in the niche through a large number of matching ligand-receptor pairs, including CSF1 expressed by dNKs, and CSF1 receptor (CSF1R) expressed on dMafs (**Extended Data fig. 4d**).

**Fig. 4.**
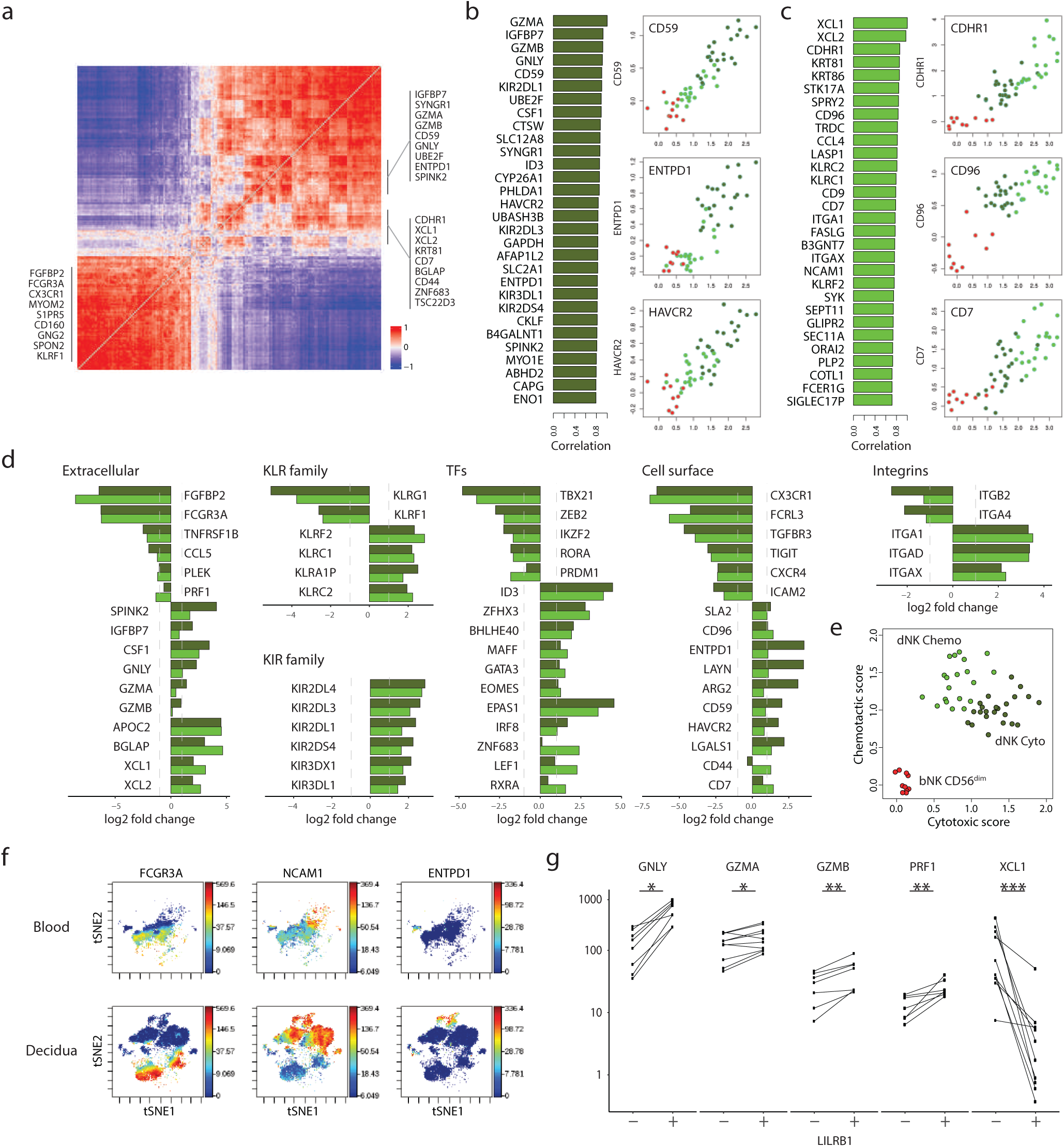
Molecular characterization of human decidual NK cells. **a,** Heatmap showing gene-gene correlation of top variable genes within NK metacells with gene modules from three distinct NK subsets. **b**, Barplot showing the top 30 genes that are most correlated with *GZMA* across NK metacells. This set of 30 cytotoxic genes is subsequently used to calculate cytotoxic scores. Scatterplots depicting the cytotoxic score per metacell versus log enrichment of a selected set of chemotactic genes. **c**, Similar to panel **b** but showing the top 30 genes most correlated with *XCL1*, defining the chemotactic score, and scatterplots depicting per metacell the chemotactic score versus log enrichment of several cytotoxic genes. **d**, Barplots showing expression fold change (log2 scale) of selected genes in human cytotoxic and chemotactic decidual NK cells when compared with circulating blood NK cells. Genes are grouped into gene families. **e**, Scatterplot depicting cytotoxic score versus chemotactic score in all NK metacells. **f**, Heatmap showing expression levels from Mass Cytometry time-of-flight (CyTOF) assay of selected proteins (NCAM1 (CD56), FCGR3A (CD16) and ENTPD1 (CD39)) on a tSNE map of all analyzed CD56+, CD3-cells from healthy donor peripheral blood and representative decidua sample. **g**, Paired point plot showing intracellular protein levels measured by flow cytometry in cytotoxic-dNKs (LILRB1^+^) and chemotactic-dNKs (LILRB1^-^).

### 3. Comparison between human and mouse immune landscape during first trimester

In order to determine the common and divergent immune populations in mouse and humans, we compared the single-cell data from the two species using a similarity-based approach^50^ (**see methods**). This analysis enabled us to project the cell populations regulating the decidual niche in the two species into a common manifold (**Fig. 3a**). Many immune populations were shared between human and mouse early to mid-pregnancy (**Fig. 3b**). In general, the mouse MFI was characterized by higher proportion of myeloid cells as compared to the human MFI, which contained mostly lymphoid populations **(Fig. 3c)**. Among the common cell states, we detected high conservation in the T and NK groups. Decidual T cells in both human and mouse displayed similar expression programs, including genes such as CXCR6, CCR5 and genes related to T cell dysfunction e.g. *PDCD1* (PD-1), *LAG3*, and *CTLA4* expressed by dT (**Fig. 3d**).

### 4. Decidual NK heterogeneity

The main decidual lymphoid cells we observed were the CD56^bright^CD16^-^ NK cells (dNK). In early first trimester, dNKs constitute 80% of all immune cells in the human decidua. dNKs in both species expressed dysfunctional and hypoxia markers, such as *HAVCR2* (*Tim-3*), *ITGA1*, *PHLDA1*, and *EPAS1* (Hif-2α) (**Fig. 3e**). This may suggest that microenvironment signals such as hypoxia might direct dNK towards a more inhibitory state through the expression of *HAVCR2*, *LGALS1* and other immune modulatory genes to promote maternal tolerance. Further analysis of dNK metacells revealed three gene modules, indicating heterogeneity within the dNK group as proposed in a recent study^28^ (**Fig. 4a and Extended Data Fig. 5a**). The three modules included a first population (Dysfunctional dNK) characterized by high expression of Havcr2 (TIM-3), ENTPD1, LAYN and several cytotoxic granules (23.7 ± 17.0%), a second population (Chemotactic dNK) characterized by high expression of chemokines and growth factors (14.2 ± 13.0%), and a third *CXCR4*^+^ dNK population (1.6 ± 1.4% of total dNKs). The latter resembled peripheral blood CD56^bright^ NK cells, and was characterized by a gene module containing *CXCR4*, *CD160*, *ITGAE*, *CCL5*, and *TIGIT* (**Extended Data Fig. 5b**). As CXCR4 is known to facilitate the recruitment of CD56^bright^ NK cells in the MFI^51^, this group may represent newly recruited dNK cells.

### 5. LILRB1 expression defines two functional dNK subgroups

The second subset of dNKs (chemo-dNK) expressed *CDHR1* and high levels of the chemotactic molecules *XCL1*, *XCL2*, and *CCL4* (**Fig. 4b-d**). The chemokines XCL1/XCL2 act by binding to the receptor XCR1, which is expressed mainly on DCs (**Extended Data Fig. 4d**) ^52,53^ and to some extent on trophoblasts^54^. In addition, XCL1-XCR1 interaction has been shown to promote trophoblast invasion^54^. Our finding that decidual DCs express immunomodulatory genes such as *IDO1* and *LGALS1* **(Fig. 2e)** may suggest that in the context of the decidua, dNK-DC interactions regulate the maternal immune response towards tolerance^55,56^. Interestingly, the chemotactic dNKs expressed higher levels of *LEF1* and were defined by the unique expression of *ZNF683* (HOBIT) (**Fig. 4d**). HOBIT expression was shown to be important for tissue retention and this might link the Chemo dNK to be tissue resident).

Our data, in line with recent works^57^, propose that both dNK phenotypes differ substantially from peripheral blood NK cells (**Fig. 4e**), which are characterized by the expression of *TBX21* (T-bet), *ZEB2*, and *IKZF2* (**Fig. 4d**). We further confirmed the segregation of dNKs into subpopulation by mass cytometry by time-of -flight (CyTOF) analysis. When compared to peripheral blood NK cells, the dNKs (NCAM^+^, FCGR3A^-^) subgroups displayed differential levels of ENTPD1, ITGAE and KIRs, in agreement with the transcriptional data (**Fig. 4f and Extended Data Fig. 5c-d**).

The dysfunctional dNK (Dys-dNK) subset was characterized by expression of *HAVCR2*, *ENTPD1* and *CSF1* as well as several cytotoxic granule genes including *GNLY*, *GZMA*, *GZMB*, and *GZMK* (**Fig. 4b, 4d**). The expression of cytotoxic molecules by the Dys-dNK cells suggests cytotoxicity properties, however, considering the dysfunctional state of dys-dNK cells and the poor killing ability of uterine NK cells, they are unlikely to be effective killers. Interestingly, dys-dNK showed high expression of *LILRB1* (**Extended Data Fig. 5e**), a receptor for all HLA molecules^58^ including the non-classical HLA-G expressed selectively by trophoblasts^59^. In a recent study, LILRB1 was shown to be co-expressed with NKG2C (KLRC2) on dNKs and to be overexpressed in subsequent pregnancies. These pregnancy-trained dNKs (PTdNKs) produced higher levels of IFNG and VEGFA in vitro^60^. NK cell cytotoxicity in the decidua is important in anti-viral responses, but may jeopardize the semiallogeneic fetus. Human dys-dNKs, like their murine equivalents, express high levels of the co-inhibitory receptor *HAVCR2* (*TIM-3*). Interestingly, *LGALS9*, one of the ligands for TIM-3, is expressed by immune and non-immune cells in the MFI^61,62^. Other components of immune tolerance and regulation mechanisms were also expressed on dys-dNKs, including *ENTPD1* (*CD39*), *LAYN*, *ARG2* and *LGALS1* (**Fig. 4b-d**), as well as several cytokines and transcription factors involved in dysfunctional CD8 T cells, such as CSF1 and ID3. The dys-dNK population also expressed higher levels of both inhibitory and activation killer-cell immunoglobulin-like receptors (KIRs)^63,64^ (**Fig. 4b-d**). This observation may suggest that they are involved in cell killing, as it was shown that higher KIR expression correlates with NK cytotoxicity^65^. To test this hypothesis, we performed several functional assays comparing the cell killing activity of chemotactic and dys-dNKs. We isolated the two NK cell populations based on the expression of LILRB1 as a marker. Using flow cytometry, we confirmed that the dys-dNKs (LILRB1^+^) expressed high protein levels of the killing factors GZMA, GZMB, GNLY, PRF1 and that the chemotactic dNKs (LILRB1^-^) expressed higher protein levels of XCL1 (**Fig. 4g**), consistent with the transcriptomic data. We then measured the killing efficiency of the dys-dNKs and the chemotactic dNKs by a killing assay and observed that the dys-dNKs were significantly more efficient in target cell killing (**Extended Data Fig. 5f**).

### 6. Tumor and decidual NK cells display overlapping characteristics

Malignancies may exploit immune modulatory mechanisms that support pregnancy. Under this premise, we asked whether the different dNK subtypes found in the MFI are also present in the TME. To explore this possibility, we mined human cancer scRNA-seq data repositories - Tumor Immune Cell Atlas for precision oncology^70^ and Wu et al.^71^. We used marker genes to purify Tumor Associated NK cells (TANKs) from pre-annotated lymphocytes. This resulted in 8,963 TANKs from 13 cancer subtypes (only 10X chromium generated data was analyzed) that were jointly subjected to MetaCell^36^ analysis.

Metacells were further clustered, allowing the identification of pan-cancer TANK subsets with a high correspondence to the three previously-defined major dNK subsets, i.e., peripheral blood TANKs (*KLRF1, FGFBP2, SPON2, ITGB2, FCGR3A*), dysfunctional TANKs (*GNLY, GZMA, ENTPD1, ITGA1, KLRC1, “*dys-TANKs”) and chemotactic TANKs (*XCL1, XCL2, IL7R*) (**Fig 5.a, Extended Fig. 6.a-b**). The expression of *IL7R* by the Chemotactic TANKs suggest they acquire a non-cytotoxic, “ILC1-like” phenotype in the TME as previously suggested^72^. We hypothesize that dys-TANKs subsets within the TME of patients might be a tumor type-specific phenomenon, driven by a unique milieu of the TME. Indeed, when focusing on three types of breast cancer cohorts, we found that the dys-TNAKs are mostly present in triple negative breast cancer tumors **(Extended Fig. 6c)**, which is the most aggressive type of breast cancer.

**Figure 5.**
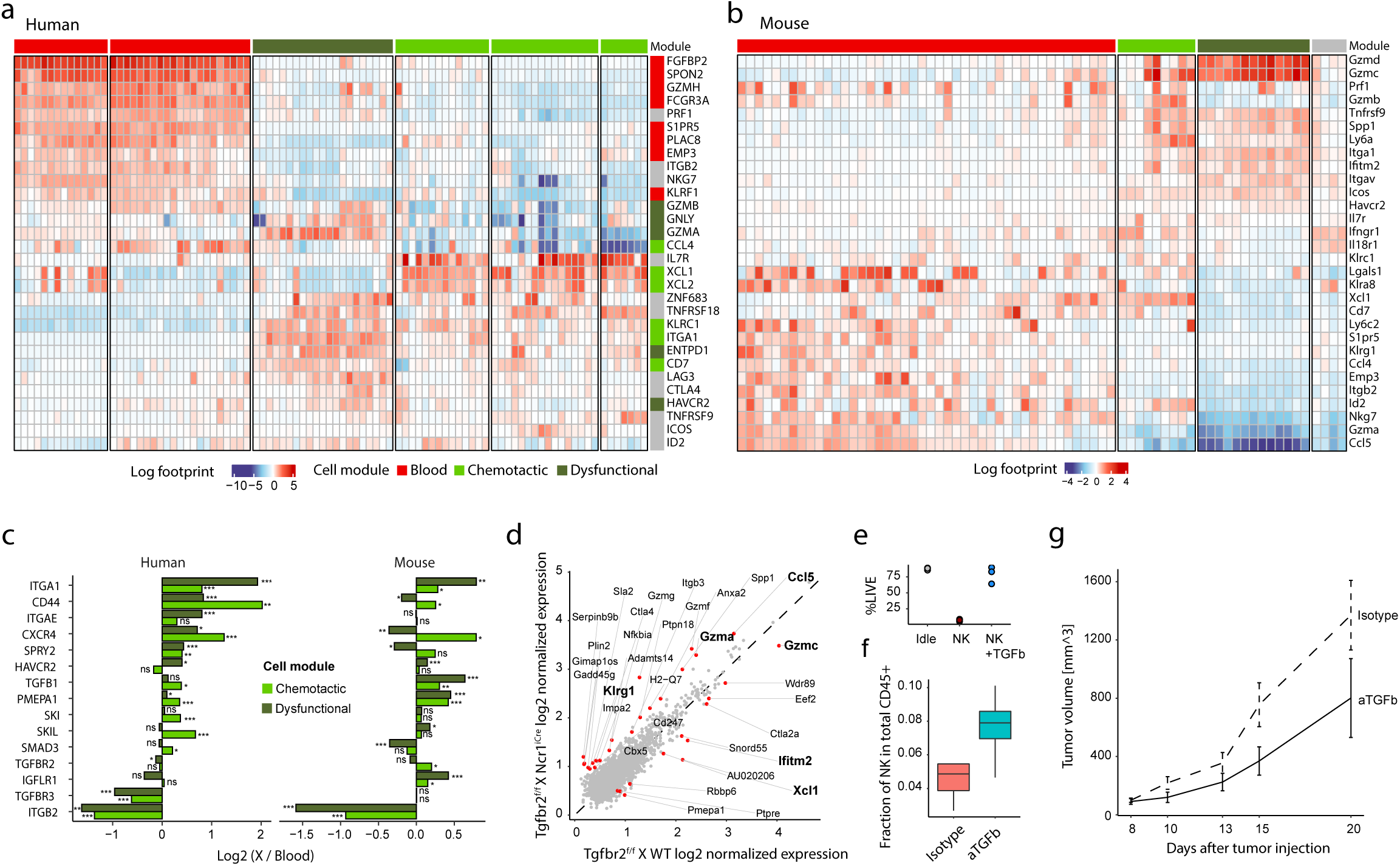
TGFbeta drives NK dysfunction in human and mouse tumors. **a**, Heatmap of human NK cells derived from diverse tumors. Columns represent metacells, further aggregated into 6 clustered representing 3 distinct modules. Color-coded for log scaled gene expression. **b**, Same as a but for murine NK cells. **c**, Barplots showing expression fold change (log2 scale) of selected genes in cytotoxic and chemotactic TANKs compared with blood TANKs. **d**, Scaled gene expression of TGFb KO vs WT NK cells. **e**, Live fraction of K562 cells post incubation with NK cells, NK cells + TGFb, or idle. **f**, Fraction of NK cells from total CD45+ cells in anti-TGFb-treated and isotype control tumors. **g**, Tumor kinetics for mice treated with anti-TGFb and isotype control.

Next, in order to test if the three NK subtypes are also present in the murine TME, we generated a single-cell atlas of NK cells isolated from peripheral blood and several mouse syngeneic tumor models: B16F10 melanoma, MC-38 colorectal, MCA-205 fibrosarcoma and Eo771 breast cancer. NK cells were sorted based on the expression of NK1.1 and Ncr1 (CD3 negative). Clustering reveals four well-defined NK cells signatures **(Fig 5b, Extended Fig. 6d)** three of which consistent with human signatures, namely blood TANKs (*Emp3, Ccl4, Itgb2* and *Klra8*), chemotactic TANKs (*Xcl1* and *Ly6a*), and dysfunctional TANKs (*Itgav, Itga1, Gzmd* and *Gzme*).

We next asked what signals in the TME may drive the unique signature of the two TANKs populations. Dissecting the NK subsets by functional genes, we find several genes that act downstream of TGF-β, e.g., *ITGA1* and *PMEPA1*, to be upregulated in non-blood TANKs **(Fig. 5c)** in both species. This suggests TGF-β may drive these TANK phenotypes. To verify this, we utilized a mouse model permitting NK specific deletion of the TGFBRII gene. This was achieved by crossing NCR1^cre^ with TGFBRII^flox^ mice. We then challenged these mice, along with WT X TGFBRII^flox^ controls, with MCA-1956 sarcoma model and sorted general immune (CD45^+^) and NK cells (CD3^-^, NK1.1^+^) at day 13 post tumor inoculation and performed scRNA-seq (**Extended Fig. 6e**). Following clustering and annotation (**Extended Fig. 6f-g**), we find that, in agreement with our hypothesis, NCR1^cre^ derived NK cells are enriched for blood NK genes (*Ccl5*, *Gzma*), compared to TANKs genes (*Xcl1, Gzmc, and Ifitm2*) in WT (**Fig. 5e**).

Finally, having identified TGF-β as the upstream effector in the tumor context, we performed in vitro and in vivo functional assays to evaluate its effect on tumor cell killing. K562 cells were co-cultured with murine NK cells in the presence or absence of TGF-β cytokine. In line with the TGF-β driven changes in transcriptional signatures towards a less effector NK cell state, the killing abilities of NK cells were dramatically reduced in the TGF-β+ co-culture (**Fig. 5e)**, further supporting the inhibitory effect of TGF-β on NK cells function in the TME. In an in vivo setting, WT mice treated with anti-TGF-β showed increased tumor NK cell infiltration (**Fig. 5f**) and diminished tumor growth (**Fig. 5g**) compared to isotype controls.

Taken together, our data indicate that TGF-β serves as a master regulator of NK cell phenotype and activity in the TME. In-depth characterization of the NK compartment implicates TGF-β signaling as a driver of NK cell inhibition and downregulation of functional molecule, supporting the potential targeting of the TGF-β pathway in NK-cell-based therapy.

## Discussion

The immune compartment of the maternal-fetal interface (MFI) presents a unique and dynamic microenvironment where the immune system must balance between promoting fetal tolerance and ensuring maternal and fetal protection. This study offers a comprehensive characterization of the immune landscape within the MFI of both humans and mice, uncovering significant insights into the cellular dynamics and molecular pathways that govern this balance. Our findings reveal a high degree of conservation in the T and NK cell compartments across species, highlighting the fundamental role of these immune cells in maintaining the delicate equilibrium at the MFI.

One of the most striking observations in our study is the identification of distinct NK cell subpopulations within the decidua. These subpopulations exhibit divergent transcriptional profiles and functional specializations. The dysfunctional decidual NK (dys-dNK) subset, characterized by high expression of immune inhibitory receptors such as LILRB1 and HAVCR2 (TIM-3), appears to play a role in fetal protection. This subset’s interaction with trophoblast cells, facilitated through LILRB1, suggests a mechanism for controlling trophoblast invasion and ensuring fetal survival. This protective role is further supported by the expression of cytotoxic granules and other immune modulatory genes, which may help in pathogen defense while maintaining tolerance to fetal antigens.

In contrast, the chemotactic dNK subset expresses high levels of chemokines and growth factors, indicating a role in recruiting and orchestrating various immune and non-immune cell populations within the MFI. This recruitment is essential for creating a supportive microenvironment for fetal development. The identification of CXCR4+ dNK cells, which resemble peripheral blood CD56bright NK cells, suggests ongoing recruitment and tissue residency dynamics at the MFI, further emphasizing the complexity and adaptability of the immune compartment during pregnancy.

Our data also underscore the functional plasticity of decidual macrophages (dMafs) and dendritic cells (DCs). dMafs display anti-inflammatory and tissue repair functions, which are critical for maintaining a supportive environment for fetal growth. The identification of two major DC subtypes, cDC1 and cDC2, along with pDCs, highlights the diverse antigen-presenting capabilities and immune regulatory functions present at the MFI. The reduced migratory capacity and altered antigen presentation observed in decidual DCs suggest a finely tuned mechanism to prevent maternal immune activation against fetal antigens.

The parallels between the immune regulatory mechanisms at the MFI and those observed in the tumor microenvironment (TME) are particularly noteworthy. Both settings involve immune modulation, angiogenesis, hypoxia, and neo-antigen presentation, which drive immune compartments towards an inhibitory state. Recent studies have further elucidated the complexity of NK cell heterogeneity within the TME, revealing distinct NK cell subpopulations with varying degrees of cytotoxic potential and regulatory functions, analogous to the diversity observed within the MFI^73,74^. Our findings that NK cells in both MFI and TME exhibit similar dysfunctional programs, driven by TGF-β signaling, provide compelling evidence of shared immune evasion strategies. In the context of cancer, the dys-TANK subset displays a transcriptional profile akin to the dys-dNKs, suggesting that tumors may exploit these immune modulatory pathways to escape immune surveillance.

The role of TGF-β as a master regulator of NK cell dysfunction is further substantiated by our functional assays. We demonstrated that blocking TGF-β signaling in vivo enhances NK cell infiltration and anti-tumor activity, offering a potential therapeutic avenue for cancer treatment. We also showed that TGF-β cytokine is an inhibitor of NK cell tumor-killing potency in vitro. This finding aligns with previous studies that identified TGF-β as a key suppressor of NK cell function in various tissues and pathologies. By targeting this pathway, we can potentially remodel the NK cell compartment to unleash its full anti-tumor potential.

In conclusion, our comprehensive characterization of the immune landscape at the MFI offers valuable insights into the cellular and molecular mechanisms that balance fetal tolerance with immune protection. The parallels with tumor immune evasion strategies underscore the broader relevance of these findings. By elucidating the role of key regulatory pathways, such as TGF-β signaling, our study paves the way for developing novel immunotherapeutic strategies that leverage the unique properties of NK cells. Future research should focus on further dissecting the interactions between different immune cell subsets and their microenvironmental cues to fully understand their contributions to maternal-fetal health and disease.

## Methods

### Mice

Myr-Venus^66^ (YFP expressing) mice were kindly provided by Jacob Hanna lab (Weizmann Institute of Science, Rehovot, Israel). *Ifnar*^-/-^ KO mice^67^ were kindly provided by Michal Schwartz (Weizmann Institute of Science, Rehovot, Israel) and Marco Printz labs (Institute of neuropathology, University Medical Centre Freiburg, Germany). WT C57BL/6 and BALB/c mice were purchased from Harlan. All animals were handled according to the regulations formulated by the Institutional Animal Care and Use Committee (IACUC). We established a model of transgenic mice reporter system by the breeding of C57BL/6 females with YFP Myr-Venus males, in which we can discriminate between the maternal and the fetal immune responses within the placenta. Mice were housed under specific-pathogen-free conditions at the Animal Breeding Center of the Weizmann Institute of Science.

### Isolation of leukocytes from murine placenta and decidua

Female mice (8-12 weeks) were mated with Myr-Venus (YFP expressing) or with Balb/c males. The day when a vaginal plug was detected in a timed mating was counted as embryonic day 0.5, and pregnant females were sacrificed at three different time points (E8.5, E10.5, and E14.5). We isolated maternal immune cells (CD45+,YFP-) from the maternal fetal interface (placenta and decidua tissues) as previously described^68^, briefly, for tissue dissociation we performed mechanical (using scissors) and enzymatic digestion. tissues were suspended in 1ml Accutase (Sigma-Aldrich) and incubated for 15 minutes at 37°C. Next, cells were filtered through a 100-µm cell strainer, washed with ice cold sorting buffer (PBS supplemented with 0.2mM EDTA pH8 and 0.5% BSA), centrifuged (5 min, 4°C, 300g), and resuspended in ice cold sorting buffer.

### Isolation of human decidual leukocytes

Human specimens of decidua were obtained from elective first trimester (6-14 weeks) termination of normal pregnancies. Experiments were performed according to the principles of the Helsinki Declaration (0423-10-HMO), and following consentance agreements obtained from all women. An ultrasound was performed prior to all elective terminations to estimate fetus age. Any fetal anomalies with known or suspected placental disorders or malformations were not included.

Decidual tissues were rinsed of blood clots and trimmed to 1mm pieces in PBS supplemented with 1mM of penicillin streptomycin. Tissue pieces were then suspended with 15ml warm RPMI-1640 supplemented with DNase type I (0.1mg/ml; Roche) and Collagenase type IV (1mg/ml; Worthington). Enzymatic digestion was performed with frequent agitation at 37°C for 15 min. Cells were then passed through a 40µm cell strainer, washed twice with RPMI-1640 and subjected to sterile density gradient separation by density centrifugation media (Ficoll-Paque; GE Healthcare Life Sciences) in a 1:1. Centrifugation (460 g, 25 min,) was performed at 10 °C, and the mononuclear cells were carefully aspirated and washed with ice-cold FACS buffer.

### Peripheral blood of healthy women first trimester pregnancy and Isolation of peripheral blood PBMC

Blood was taken from 5 decidua donors. The peripheral blood collecting samples is part of the 0423-10-HMO approval. PBMCs were purified from fresh blood samples, derived from 5 pregnant donors, on a Ficoll gradient as previously described. Followed by red blood lysis (Sigma-Aldrich) for 5min at 4°C and washing with ice cold FACS buffer.

### Flow cytometry

Cell populations were sorted with FACSAria II SORP (BD Biosciences, San Jose, CA), FACSAria III (BD Biosciences, San Jose, CA), or with FACSAria Fuison instrument (BD Biosience, San Jose, CA).

Mouse samples were stained using the following antibodies:

CD45 (APC, eBioscience, clone: 30-F11, cat. No. 17-0451-82), Ter-119 (EF450, eBioscience, clone: TER-119, cat. No. 48-5921-82), Ly6g (EF450, eBioscience, clone: 1A8, cat. No. 48-9668-82), Ncr1 (PE, Biolegend, clone: 29A1.4, cat. No. 137603), TCR-b (FITC, Biolegend, clone: H57-597, cat. NO. 109205), CD4 (APC/CY7, Biolegend, clone: GK1.5, cat. No. 100414), CD25 (PE, Biolegend. Clone: PC61, cat. No. 102007), Foxp3 (AF488, Biolegend, clone: 150D, cat. No. 320011).

Human samples were stained using the following antibodies:

CD45 ((PerCP-Cy5.5, Biolegend, clone EO1, cat. no. 368504), CD56 (Cytognos, clone: C5.9, cat. no. CYT-56PE), CD16 (PE/cy7, Biolegend, clone: 3gG8, cat. no. 302015), CD16 (BV510, Biolegend, clone: 3gG8, cat. No. 302047), CD3 (FITC, Beckman Coulter, cat. No. A07746), CD19 (Beckman Coulter, clone: J3.119, cat. no. IM3628U), CD8 (APC-H7, BD Bioscience, clone: SK1, cat. No. 560179), CD14 (APC-Cy7, BD Bioscience, clone: MOPg, cat. No. 557831), LILRB1 (FITC, R&D systems, clone:292305, cat. No. FAB-20171F), CD96 (PE/cy7, Biolegend, clone: NK92.39 cat. no. 338415), CD7 (APC, Biolegend, clone: CD7-6B7, cat. No.343107), CD44 (APC/cy7, Biolegend, clone: IM7, cat. No. 103027), XCL1 (APC, LSbio, cat. no. LS-C716131-100), PRF1 (APC, Biolegend, clone: gD9, cat. No. 308112), GZMB (AF647, Biolegend, clone: GB11, cat. No. 515406), GZMA (APC, LSbio, cat. no. LS-C715841-100), GNLY (AF647, Biolegend, clone: DH2, cat. no. 348006).

### Intracellular staining

Intracellular staining for human NK cells was done using the Cytofix-Cytoperm Intracellular Staining kit (BD Bioscience), according to the manufacturer’s instructions. Membrane staining (CD56, CD3, CD16, LILRB1) was followed by intracellular staining of various intracellular factors (Xcl1, Gzma, Gzmb, Prf1 and Gnly). All samples were analyzed by flow cytometry software (Kaluza Analysis software and FCS Express 4), while analyzing separately the intracellular staining on the CD56^+^CD3^-^ CD16^-^LILRB1^+^ and CD56^+^CD3^-^CD16^-^LILRB1^-^.

### Mass Cytometry (Cytof)

Frozen samples from pregnant donors and healthy controls were thawed and stained for Cytof. The Cytof panel included the listed antibodies purchased from Fluidigm: CD45-089Y (3089003B), CD196-141Pr (3141003A), CD19-142Nd (3142001B), CD117-143Nd (3143001B), CD4-145Nd (3145001B),CD8a-146Nd (3146001B), CD25-149Sm (3149010B), CD103-151Eu (3151011B), TCRgd-152Sm (3152008B), TIGIT-153Eu (3153019B), CD3-154Sm (3154003B), CD194-158Gd (3158032A), CD161-164Dy (3164009B), CD39-160Gd (3160004B), CD27-162Dy (3162009B), CD45RO-165Ho (3165011B), CD127-168Er (3168017B), CD159a-169Tm (3169013B), CD45RA-170Er (3170010B), CD226-171Yb (3171013B); CD94-174Yb (3174015B), CD14-175Lu (3175015B), CD56-176Yb (3176009B); CD16 −299Bi (3209002B), CD7-147Sm (3147006B), PD1-155Gd (3155009B), CD26-161Dy (3161015B), CD57-163Dy (3163022B), CD158b-173Yb (3173010B), ICOS-148Nd (3148019B), CXCR3-156Gd (3156004B), CCR7-159Tb (3159003A), FceRI and CD138-150Nd (3150027B; 3150012B). NKp44-167Er, which was conjugated in-house (eBioscience, 16-3369-85), was also included in the panel.

Cells were washed with Cy-FACS buffer (CyPBS, Rockland, MB-008; 0.1% BSA, Sigma, A3059; 0.02% sodium azide, Sigma, 71289, 2 mM ethylenediaminetetraacetic acid, Hoefer, GR123-100) and stained on ice for 1 h. After two washes, cells were stained with cisplatin (Enzo Life Sciences, NC0503617) for 1 min, washed again twice, fixed in 4% paraformaldehyde (Electron Microscopy Sciences, 15710) for 15 min, spun down and re-suspended in Intercalator–Ir125 (Fluidigm, 201192A) overnight. Cells were washed and analyzed on a Cytof 2 mass cytometer (Fluidigm). Data was processed using Cytobank.

### Single cell sorting

Prior to sorting, samples were filtered through a 70 μm mesh. For the sorting of whole immune cell populations, samples were gated for CD45^+^.

For sorting of human decidual NK cells samples were gated for CD45^+^, CD3^-^, CD56^+^ (“bright”). For sorting of human non-decidual NK cells samples were gated for CD45^+^, CD56. For sorting of murine maternal immune cells, cells were gated on CD45^+^, YFP^-^, Ly6g^-^. Isolated live cells were single cell sorted into 384-well cell capture plates containing 2 μL of lysis solution and barcoded poly(T) reverse-transcription (RT) primers for single-cell RNA-seq^69,35^. Four empty wells were kept in each 384-well plate as a no-cell control during data analysis. Immediately after sorting, each plate was spun down to ensure cell immersion into the lysis solution, and stored at −80°C until processed.

### MARS-seq 2.0 library preparation

Single-cell libraries were prepared as previously described^69,35^. In brief, mRNA from cell sorted into cell capture plates were barcoded and converted into cDNA and pooled using and automated pipeline. The pooled sample is then linearly amplified by T7 *in vitro* transcription, and the resulting RNA is fragmented and converted into sequencing-ready library by tagging the samples with pool barcodes and illumine sequences during ligation, RT, and PCR. Each pool of cells was tested for library quality and concentration is assessed as described earlier^69,35^. Overall, barcoding was done on three levels: cell barcodes allow attribution of each sequence read to its cell of origin, thus enabling pooling; unique molecular identifiers (UMIs) allow tagging each original molecule in order to avoid amplification bias; and plate barcodes allow elimination of the batch effect.

### Bulk RNA-seq

50,000 CD45+ or CD45-cells from tumors were sorted directly into 50 ul of lysis binding buffer (Thermo Fisher Scientific), vortexed and stored at −80c. RNA was extracted from the lysed cells by using Dynabeads™ Oligo(dT) mRNA isolation kit (Thermo Fisher Scientific). RNA next-generation sequencing was performed using a modified version of MARS-seq, as previously described^69^. TMM normalization (edgeR) was applied to read counts to account for library size and composition.

### Low-level MARS-seq processing

scRNA-seq libraries (pooled at equimolar concentration) were sequenced on an Illumina NextSeq 500 at a median sequencing depth of ∼40,000 reads per cell. Sequences were mapped to the mouse genome (mm9) or human genome (hg38) accordingly, demultiplexed, and filtered as previously described^69,35^ with the following adaptations. Mapping of reads was done using HISAT (version 0.1.6); reads with multiple mapping positions were excluded. Reads were associated with genes, if they were mapped to an exon using the annotation from UCSC genome browser as reference. Exons of different genes that shared a genomic position on the same strand were considered as a single gene with a concatenated gene symbol. We estimated the level of spurious UMIs in the data using statistics on empty MARS-seq wells, and excluded rare cases with estimated noise >5% (median estimated noise over all experiment was 2%).

### Metacell modeling for MARS-seq data from mouse MFI

To analyze the MARS-seq data from mouse MFI, we used the MetaCell package^36^ with the following specific parameters. We removed specific mitochondrial genes, immunoglobulin genes, high abundance lincRNA (Table SX), and genes linked with poorly supported transcriptional models (such as genes annotated with the prefix “Gm-”, “Rik-”, and etc.). We then filtered cells with less than 500 UMIs or total fraction of mitochondrial gene expression exceeding 50%. Gene features with high variance to mean were selected using the parameter Tvm = 0.2 and minimal total umi > 100. We excluded gene features associated with the general cell stress and type-I Interferon response (Table S6) using a clustering approach. To this end we identified first all genes with a correlation coefficient of at least 0.1 with one of the anchor genes *Fos*, *Fosb*, *Nkfbib*, *Nfkbiz*, *Isg15*, *Isg20*, *Irf7*, *Mnda*, *Mndal*, *Zfp36*, *Zfp36l1*, *Ifit1*, *Ifit2*, *Ifit3*. We then hierarchically clustered the correlation matrix between these genes (filtering genes with low coverage and computing correlation using a down-sampled UMI matrix) and selected the gene clusters that contained the above anchor genes. Erythrocytes, neutrophils, and non-immune cells were considered a contamination, and were removed before performing the final clustering.

The gene selection strategy discussed above retained a total of 3,364 gene features for the computation of the Metacell balanced similarity graph. We used K = 500 bootstrap iterations. We did not apply outlier filtering, but allowed metacell splitting by clustering the cells within each metacell and splitting it, if distinct clusters are detected. Annotation of the metacell model was done using the metacell confusion matrix and analysis of marker genes.

### Metacell modeling for MARS-seq data from human MFI

Analysis of MARS-seq data from human MFI was performed similarly to mouse MFI data, but with the following specific parameters: We removed specific mitochondrial genes, immunoglobulin genes, high abundance lincRNA (Table SX), and genes linked with poorly supported transcriptional models (such as genes annotated with the prefix “AC[0-9]”, “AP[0-9]”). We then filtered cells with less than 400 UMI. We excluded gene features associated with the cell cycle and general cell stress using a clustering approach. We identified first all genes with a correlation coefficient of at least 0.1 for one of the anchor genes *MKI67*, *TOP2A*, *STMN1*, *TUBB*, *HMGB2*, *HMGB1*, *FOS*, *ZFP36*, *FOSB*, *NKFBIA*, *JUND*, *JUN*. We then hierarchically clustered the correlation matrix between these genes (filtering genes with low coverage and computing correlation using a down-sampled UMI matrix) and selected the gene clusters that contained the above anchor genes. Non-immune are removed before doing the final clustering.

### Comparison between human and mouse immune landscape

To systematically compare the immune cells from human and mouse MFI, we quantified the similarity between the average transcriptome of the immune cell types identified in the two datasets. To match the genes in human and mouse, orthologs in the two species were obtained from Ensembl BioMart. We then focused on the genes that can best distinguish the different cell types in each species. In each dataset, we selected the top 500 most variable genes according to the metacell model, and identified in total 131 genes that were shared between the two species. We then calculated the UMI counts per cell of these genes in individual cell types in both human and mice datasets, and normalized the values within each dataset (scale the values to between 0 and 1). Finally, we created a vector of 131 elements using the normalized expression of the 131 genes as footprints for each cell type in human and mouse, and computed the cosine similarity between the vectors to estimate the similarity between the different cell types in the two species.

### Defining gene modules in various human MFI NK phenotypes

To account for the complex gene expression patterns in various human MFI NK phenotypes, we combined metacell analysis with an approach aiming at identifying quantitative gene signatures, which was described previously^50^. Given any list of signature genes for a certain phenotype, we defined the signature’s scores for each metacell by averaging the metacell log enrichment scores (lfp values) of the genes in the set. Note that using this approach we limited the contribution of highly expressed genes to the score, and that we relied on the regularization of the metacell computation of gene enrichment scores to restrict the noise levels inflicted over the phenotype scores.

To define signatures gene sets for cytotoxic NK cells, chemotactic NK cells, and blood NK cells, we identified groups of 30 genes that were maximally correlated to selected anchor genes (*GZMA*, *XCL1*, and *FGFBP2*), using linear correlation over metacells’ log enrichment scores. Genes associated with cell cycle, type I IFN and stress were filtered from these lists. Anchors were validated by using alternative anchor genes that yielded similar gene signatures and more systematically by testing correlation of the signature scores computed from their derived gene sets to all genes while excluding the gene itself from the score, establishing consistency and robustness (genes remained top ranking even when omitting them from the score) to anchor selection.

### Differential gene expression analysis

To compare the gene expression between two sets of cells, the total UMI counts of individual cells were down-sampled to the same value as the one used for Metacell analysis in each dataset. Next, genes with average expression level of less than 1% cells with at least one UMI in both of the two sets of cells were excluded. Then, differentially expressed genes between the two sets of cells were identified using a two-tailed Mann-Whitney U test, with the Benjamini-Hochberg procedure for multiple hypothesis correction. Finally, Genes with FDR < 0.05 and an absolute fold-change > 1.5, were considered as differentially expressed.

### Metacell modeling for human tumor NK cells

Single-cell RNA-seq data from multiple cancer types were sourced from the Tumor Immune Cell Atlas for Precision Oncology^70^ and Wu et al. ^71^. Clustering was performed on annotated lymphocytes using MetaCell, following previously established protocols. T cells were excluded based on the expression of *CD3D, CD3G*, *CD8A*, and *CD8B*. To address batch effects, batch-specific genes were removed, and the clustering process was iterated to refine the results.

### Radioactive NK killing assay

The cytolytic activity of the LILRB1^-^ and LILRB1^+^ dNK cells was assessed in 5-hour ^35^S-release assays in which effector cells were mixed with 5,000-[^35^S] methionine-labeled target cells (721.221) at different E:T ratios in U-bottomed plates. Assays were terminated by centrifugation at 1,600 rpm for 5 min, and 50μl of the supernatant was collected for liquid scintillation counting. Percent of specific lysis was calculated as follows: % lysis = [(experimental well - spontaneous release) / (maximal release - spontaneous release)] X 100. Spontaneous release was determined by incubation of the labeled target cells with medium only. Maximal release was determined by solubilizing target cells in 0.1 M NaOH.

### NK killing with TGFb

K562 cells (1 × 10^6^) were cultured under three conditions: (1) alone (control), (2) with 100,000 NK cells isolated from PBMCs using the Miltenyi NK Cell Isolation Kit, or (3) with 100,000 NK cells in the presence of 10 µg/mL TGF-β cytokine. After 48 hours, cytotoxicity was assessed by determining the fraction of viable tumor cells, identified as DAPI-negative K562 cells.

### Tumor cell line

MCA-205 fibrosarcoma cell lines were kindly provided by Sergio Quezada group at UCL cancer institute, London, UK. MC-38 were kindly provided by Roni Dahan group from Weizmann Institute of science. B16F10, K562 and E0771 cell lines were purchased from ATCC.

Cells were cultured in DMEM (41965-039) medium supplemented with 10% heat-inactivated FBS, 1mM sodium pyruvate, 2mM l-glutamine, 1% penicillin-streptomycin (Thermo Fisher Scientific). Cells were cultured in 10cm tissue culture plates in an incubator (Thermo Fisher Scientific) with humidified air and 5% CO2 at 37°C. Cell lines were validated for lack of mycoplasma infection using primers for mycoplasma-specific 16S rRNA gene region (EZPCR Mycoplasma Kit; Biological Industries).

### Tumor growth measurements

Mice were inoculated intradermally (i.d.) with 5×10^5^ MCA-205 cells suspended in 100μl?? PBS on their right flank. At day 6, when tumors become visible, their volume was measured every two days using a caliper. Tumor volume was assessed by measuring two diameters and calculated using the formula *X*^2^ × *Y* × 0.52 (where *X*, smaller diameter and *Y*, larger diameter).

### TGFb blocking

Tumor-bearing mice were administered 100 µg of anti-TGF-β blocking antibody (BioXcell, cat: BP0057) or the corresponding isotype control every three days.

### Isolation of tumor infiltrating leukocytes

Tumor bearing mice were sacrificed at 11 or 15 days after tumor cell inoculation. The tumors underwent mechanical (gentleMACS^TM^ C tube, Miltenyi Biotec Inc., San Diego, CA) and enzymatic digestion (0.1mg/ml DNase type I; Roche, and 1mg/ml Collagenase IV; Worthington) in RPMI-1640 for 15 min at 37°C. Cells then filtered through 100µm cell strainer, washed with ice cold sorting buffer, centrifuged (5 min, 4°C, 300g), and stained with fluorophores conjugated antibodies followed by single-cell sorting.

## Acknowledgments

We thank the participants in the human clinical studies, I. Eisenberg for scientific suggestions and logistics in obtaining human samples, M. Cohen and A. Giladi for reviewing the manuscript, Genia Brodsky from the Scientific Illustration unit of the Weizmann Institute for artwork, members of the Amit laboratory for discussions, I.A. is supported by the Chan Zuckerberg Initiative (CZI), the HHMI International Scholar award, the European Research Council Consolidator Grant (ERC-COG) 724471-HemTree2.0, the Thompson Family Foundation, an MRA Established Investigator Award (509044), the Israel Science Foundation (703/15), the Ernest and Bonnie Beutler Research Program for Excellence in Genomic Medicine, the Helen and Martin Kimmel award for innovative investigation, the NeuroMac DFG/Transregional Collaborative Research Center Grant, an International Progressive MS Alliance/NMSS PA-1604-08459 and an Adelis Foundation grant, Howard Hughes Medical Institute (HHMI). I.A. is the incumbent of the Alan and Laraine Fischer Career Development Chair.

## Author contributions

A.J designed the study, performed and analyzed experiments, and wrote the manuscript. S-Y.W performed single cell RNA sequencing data analysis, analyzed experiments and wrote the manuscript. T.L performed bulk and single cell RNA sequencing data analysis and wrote the manuscript. M.G performed human samples collection and digestion, flow cytometry analysis, NK killing assays, analyzed experiments and provided input on experimental design. N.E performed single cell RNA sequencing data analysis and wrote the manuscript. D. G-W coordinated all human sample collection and provided input on experimental design. E.D performed data curation. M.M and M.C performed CyTOF experiments and analysis of human samples. R.K collected human samples and performed flow cytometry experiments. T. M-S performed flow cytometry experiments and assisted in flow cytometry analysis. R.G, R.A and D.S performed elective first trimester termination of normal pregnancies procedures and collected human samples. S.Y coordinated and supervised clinical data and provided input on experimental design. M.C and O.M provided input on experimental design and analyzed experiments. I.A supervised the project, designed and analyzed experiments and wrote the manuscript.

## Data and code availability

Will be provided upon request.

## Competing Interests

The authors declare they have no competing interests regarding this manuscript.

## Extended Data Figure Legends

**Extended Data Fig. 1.**
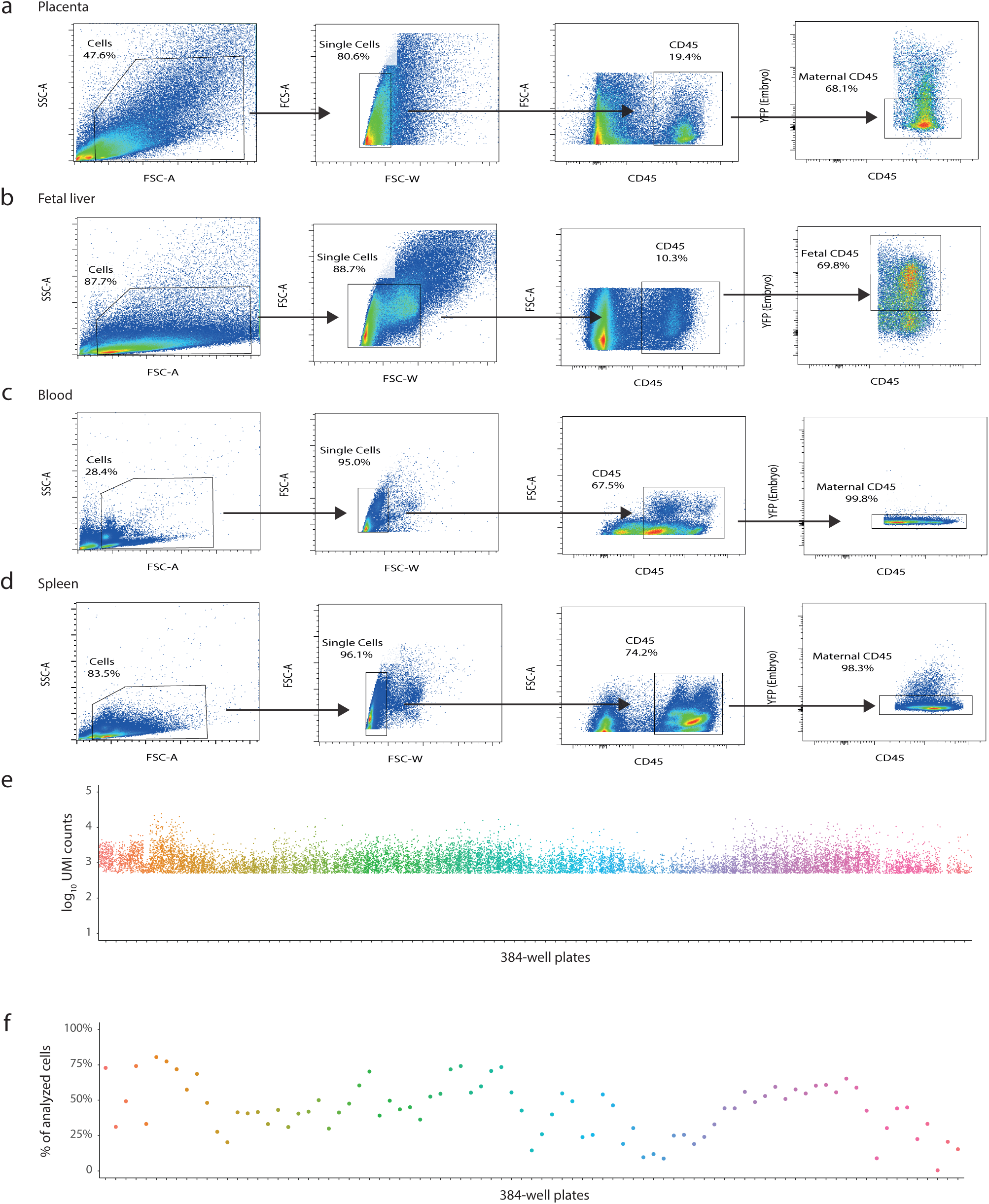
Sorting strategy and single cell RNA-seq QC values for murine maternal immune cells. **a-d**, Representative flow cytometry plots showing sorting strategy (CD45^+^, YFP^-^) maternal immune cells after doublet exclusion: **a**, maternal immune cells from E14.5 maternal fetal interface; **b**, immune cells from E14.5 fetal liver; **c**, immune cells from E14.5 pregnant female Spleen; **d**, immune cells from E14.5 pregnant female peripheral blood. **e**, Total UMIs per single cell from each sorted 384-well plate. **f**, Fraction of analyzed cells after filtering per plate.

**Extended Data Fig. 2.**
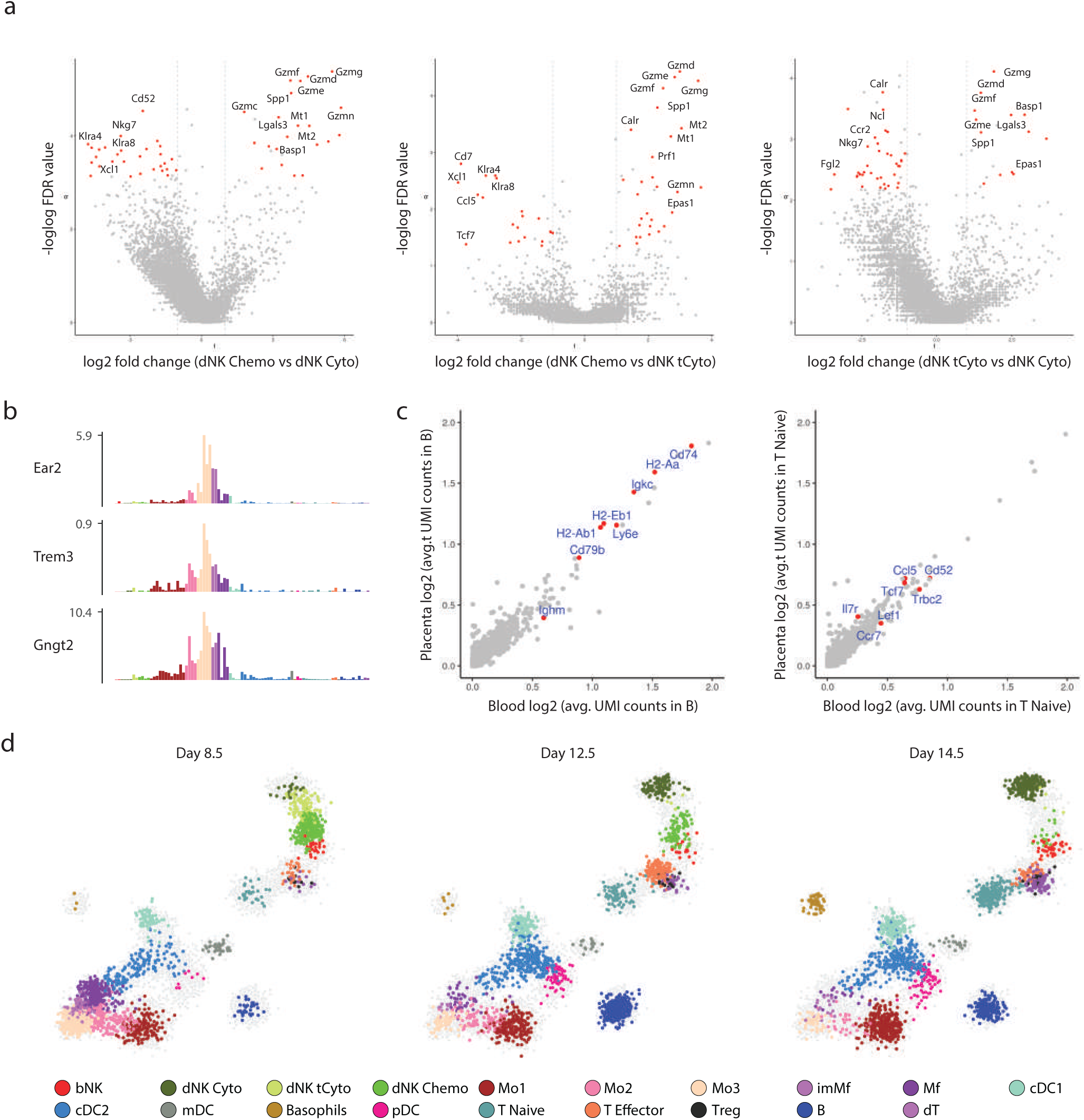
Profiling maternal immune cell landscape of mouse maternal-fetal interface across first trimester. **a,** Volcano plots showing the differentially expressed genes between mouse chemotactic decidual NK cells with mouse cytotoxic decidual NK cells, mouse chemotactic decidual NK cells with mouse transient cytotoxic decidual NK cells, and mouse transient cytotoxic decidual NK cells with mouse cytotoxic decidual NK cells, with the highlights of selected important genes. Mann-Whitney test with Bonferroni adjustment was used to calculate the FDR for the comparison. **b**, Normalized expression levels of select genes across the entire metacell model, color coded by cell type. **c**, Scatterplot on the left showing the average UMI counts (log2 scale) of B cells in blood (x axis) compared with B cells in placenta (y axis), with the highlights of selected important genes in B cells. Scatterplot on the right showing the same but for the naïve T cells in blood and naïve T cells in placenta. **d**, 2D projection of single cells taken from placenta of E8.5, E12.5, and E14.5 of pregnancy, respectively. Cells were down-sampled to 1,900 cells for each time point of the pregnancy.

**Extended Data Fig. 3.**
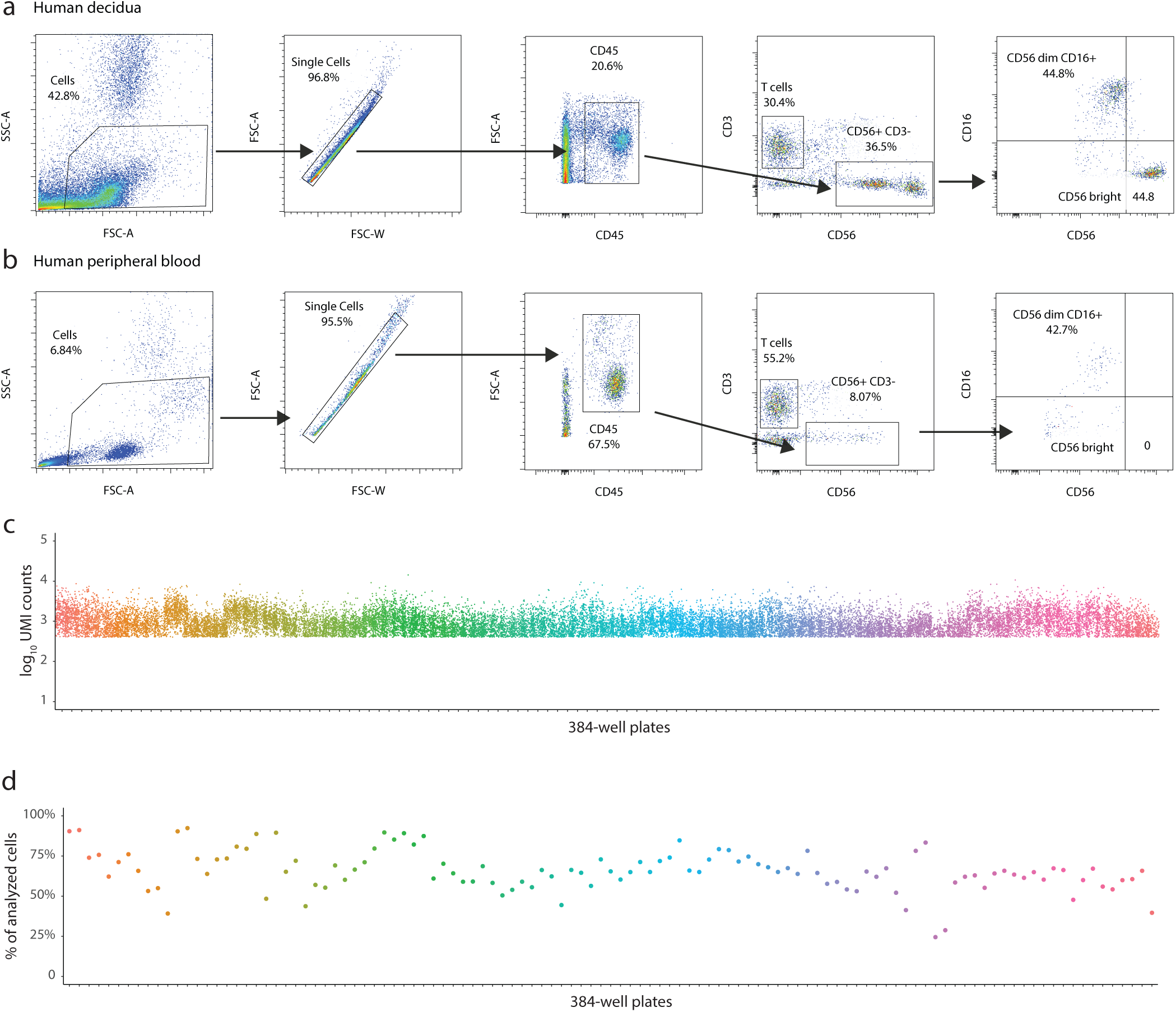
Gating strategy and single cell RNA-seq QC values for human maternal-fetal interface immune cells across first trimester pregnancy. **a-b**, Representative flow cytometry plots showing sorting strategy (CD45^+^) for immune cells or NK (CD45^+^, CD56^+^, CD3^-^) after doublet exclusion. Samples are from human decidua tissue (**a**) and peripheral blood (**b**). Cells were stained for immune populations markers such as CD45, CD3 (T cells), and CD56 (NK cells). **c**, Total UMIs per single cell from each sorted 384-well plate. **d**, Fraction of analyzed cells after filtering per plate.

**Extended Data Fig. 4.**
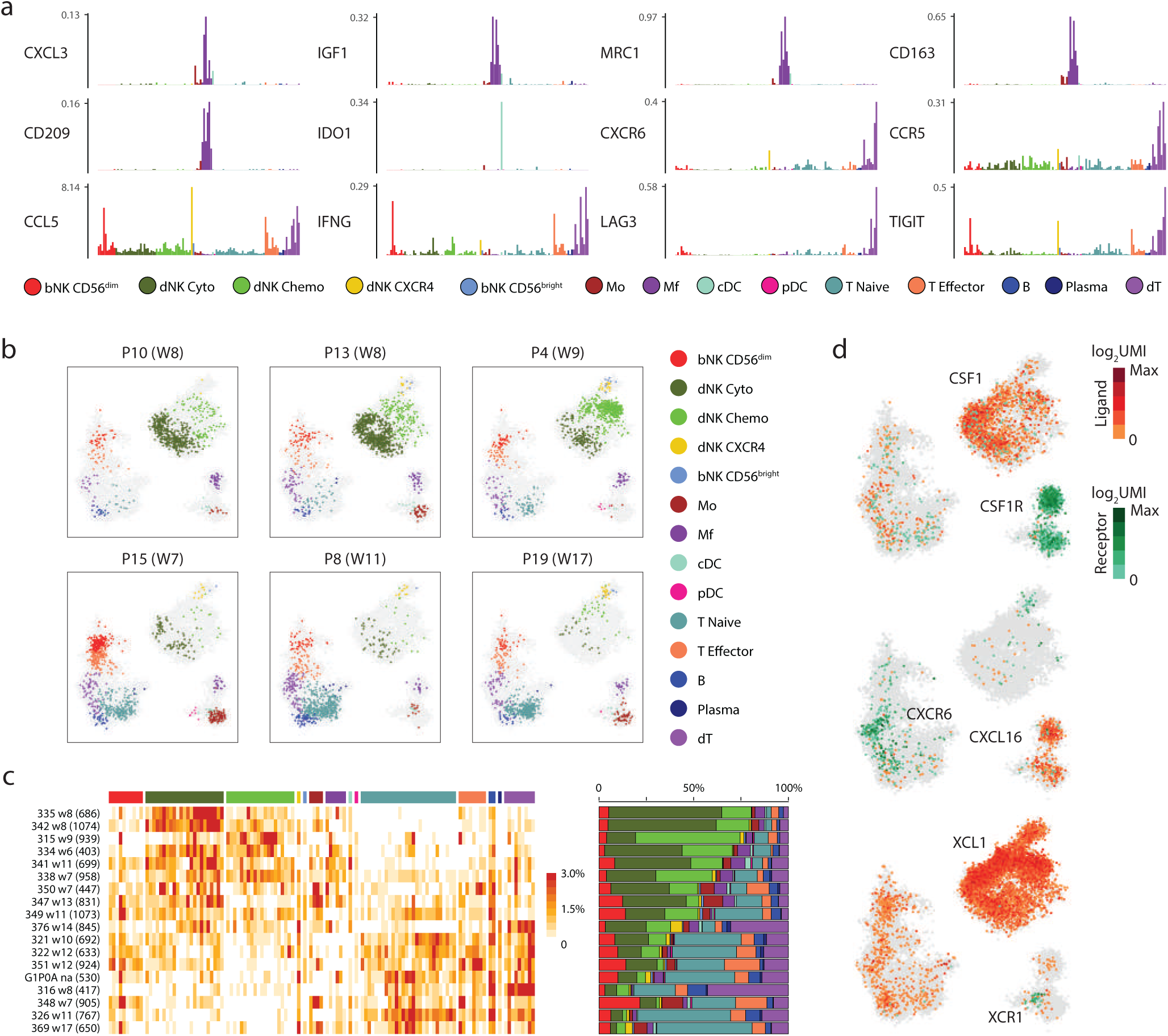
Maternal immune cell landscape of human maternal-fetal interface across first trimester. **a**, Normalized expression levels of select genes across the entire metacell model, color coded by cell type. **b**, 2D projection for single cells from three patients that are in the early weeks of the first trimester (top ones) and three patients that are in the late weeks of the first trimester (bottom ones), with individual cells shown in dots and color coded by cell type. **c**, Metacells (columns) are ordered by cell types. Patients (rows) are ordered by the frequency of decidual NK cells, with number of cells per patients in parenthesis. Compositions of different cell types are shown on the right (colors as panel **b**). **d**, Heatmap showing unique molecular identifier (UMI) counts (log2 scale) in individual cells on the 2D map of **Fig**. **2a** for selected ligand-receptor pairs. The UMI counts of a ligand are shown in red, and the UMI counts of a receptor in green.

**Extended Data Fig. 5.**
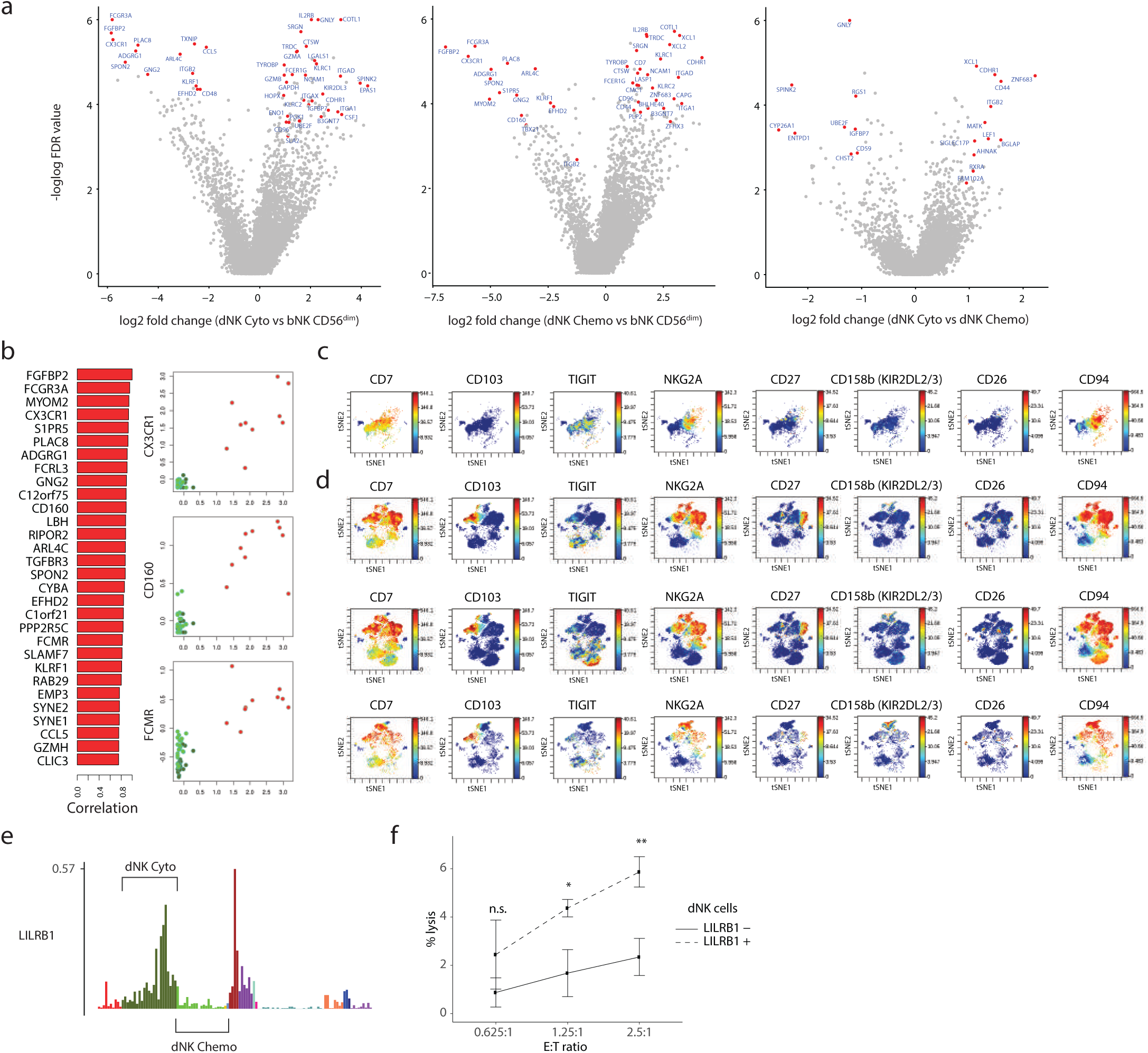
Characterization of human decidua NK cells. **a**, Volcano plots showing the differentially expressed genes between human cytotoxic decidual NK cells with human CD56^dim^ blood NK cells, human chemotactic decidual NK cells with human CD56^dim^ blood NK cells, and human cytotoxic decidual NK cells with human chemotactic decidual NK cells, with the highlights of selected important genes. Mann-Whitney test with Bonferroni adjustment was used to calculate the FDR for the comparison. **b**, Barplot showing the top 30 genes that are most correlated with *FGFBP2* across NK metacells. This set of 30 genes is subsequently used to calculate blood NK cell scores. Scatterplots depicting the blood NK cell score per metacell versus log enrichment of a selected set of blood NK cell genes. **c-d**, Mass Cytometry (CyTOF) analysis of CD56^+^, CD3^-^ cells in peripheral blood of healthy donors (**c**) and 3 decidual samples (**d**). **e**, Normalized expression levels of gene *LILRB1* across the entire metacell model, color coded by cell type, with the highlights of metacells for cytotoxic decidual NK cells and chemotactic decidual NK cells. **f.** dNK cytotoxic assay: x axis denotes effector to target cell ratio; y axis denotes mean percentage lysis of target cells by dNK cells; Stars denote p value (*p < 0.05; **p < 0.01) of a student t-test comparing assays using LILRB1^+^/LILRB1^-^ sorted dNK cell subsets.

**Extended Data Fig. 6.**
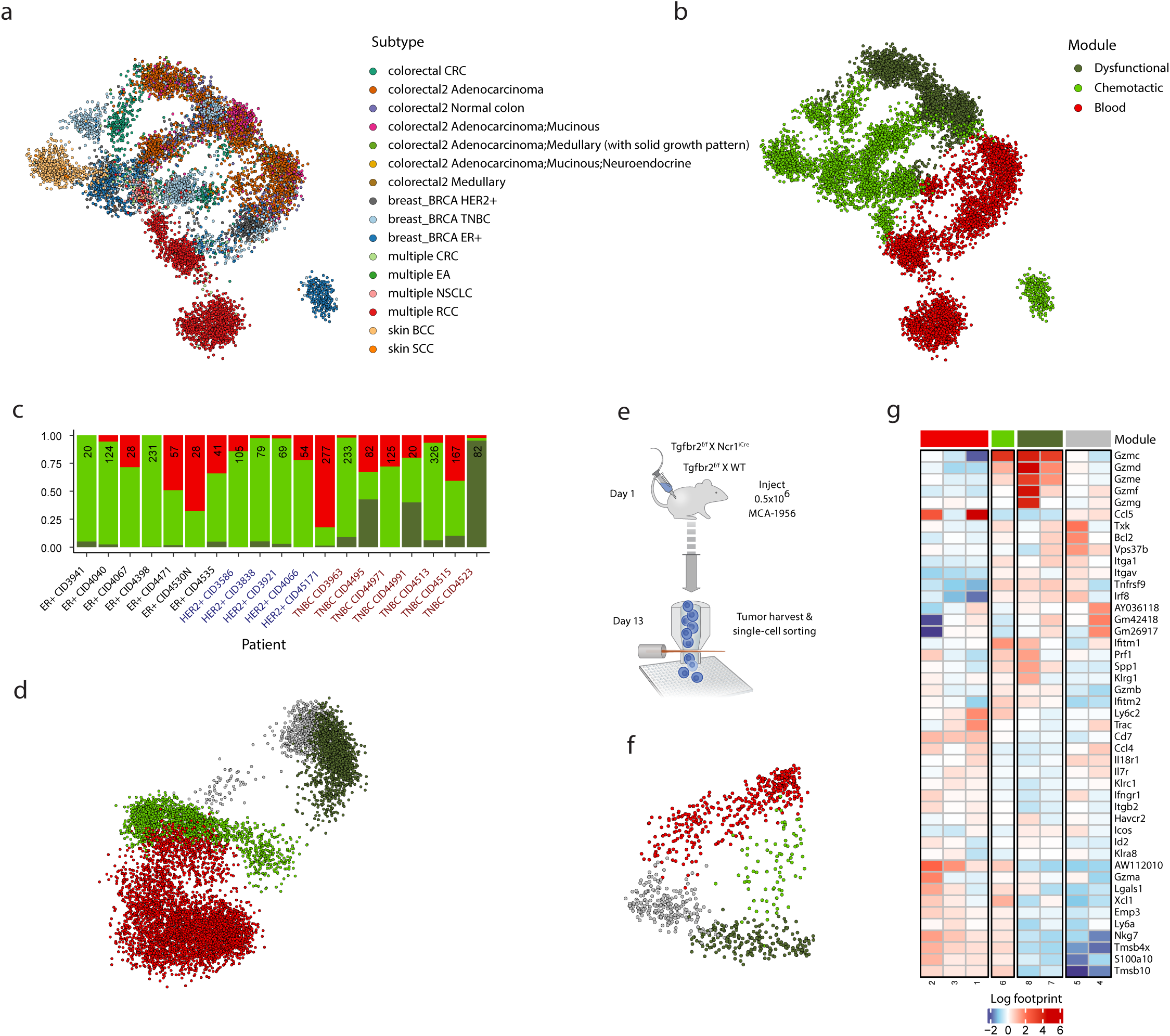
TGFbeta drives NK dysfunction in human and mouse tumors. **a**, 2D projection of transcriptome profiles of 8,963 QC-passed NK derived from 5 studies containing 13 cancer subtypes marked by color code. **b**, 2D projection of single cells clustered into three subsets marked by color code. **c**. Fraction of NK cell subsets in 3 breast cancer subtypes. **d**. 2D projection of single NK cells from diverse murine cancer types clustered into four subsets marked by color code. **e**. Experimental design to characterize the response of NK cells to Tgfbr2 KO. **f**. 2D projection of single NK cells from the model in e, clustered into four subsets marked by color code. **g**. Heatmap of murine NK cells derived from the model in (**e)**. Columns represent metacells, further clustered into 4 modules. Color-coded for log scaled gene expression.

## Supplementary Table Legends

**Table S1. Mouse maternal-fetal interface (MFI) tissues metacell data.** Related to Figure 1, 5 and Extended Data Fig.2. Footprint gene expression of 88 metacells obtained from scRNA-seq of 14,145 maternal-fetal interface tissue immune cells from wild-type and *Infar^-/-^* mice.

**Table S2. Human decidual tissue and peripheral blood metacell data.** Related to Figure 2 and Extended Data Figure 3 and 4. Footprint gene expression of 125 metacells obtained from scRNA-seq of 27,764 decidual tissue and peripheral blood immune cells from 24 healthy donors.

**Table S3. Human melanoma and breast cancer tumors and peripheral blood metacell data.** Related to Figure 4 and Extended Data Figure 6. Footprint gene expression of 365 and 128 metacells obtained from scRNA-seq of 47,704 and 18,947 T and NK cells from melanoma and breast cancer tumors and peripheral blood samples respectively

**Table S4. Bulk RNA sequencing of mouse MCA-205 tumors.** Related Extended Data Figure 7. TMM (edgeR) normalized read count of bulk RNA-seq data of ∼50,000 immune cells from 2 wild-type MCA-205 tumor bearing mice, 1 *Infar^-/-^* MCA-205 tumor bearing mouse and 2 wild-type B16F10 tumor bearing mice.

**Table S5. Mouse MCA-205 tumors metacell data.** Related to Figure 6 and Extended Data Figure 7. Footprint gene expression of 56 metacells obtained from scRNA-seq of 5,685 MCA-205 tumors immune cells from wild-type and *Infar^-/-^* mice.

## Notes

### Competing Interest Statement

The authors have declared no competing interest.

